# Comparative whole-genome analysis reveals genetic adaptation of the invasive pinewood nematode

**DOI:** 10.1101/439612

**Authors:** Jiarui Li, Xinyue Cheng, Runmao Lin, Shijun Xiao, Xinxin Yi, Zhenchuan Mao, Xi Zhang, Jian Ling, Xiaojun Kou, Xia Yan, Ji Luo, Feixue Cheng, Yilong Li, Laifa Wang, Nansheng Chen, Bingyan Xie

## Abstract

Genetic adaptation to new environments is essential for invasive species. To explore the genetic underpinnings of invasiveness of a dangerous invasive species, the pinewood nematode (PWN) *Bursaphelenchus xylophilus*, we analysed the genome-wide variations of a large cohort of 55 strains isolated from both the native and introduced regions. Comparative analysis showed abundant genetic diversity existing in the nematode, especially in the native populations. Phylogenetic relationships and principal component analysis indicate a dominant invasive population/group (DIG) existing in China and expansion beyond, with few genomic variations. Putative origin and migration paths at a global scale were traced by targeted analysis of rDNA sequences. A progressive loss of genetic diversity was observed along spread routes. We focused on variations with a low frequency allele (<50%) in the native USA population but fixation in DIG, and a total of 25,992 single nuclear polymorphisms (SNPs) were screened out. We found that a clear majority of these fixation alleles originated from standing variation. Functional annotation of these SNP-harboured genes showed that adaptation-related genes are abundant, such as genes that encode for chemoreceptors, proteases, detoxification enzymes, and proteins involved in signal transduction and in response to stresses and stimuli. Some genes under positive selection were predicted. Our results suggest that adaptability to new environments plays essentially roles in PWN invasiveness. Genetic drift, mutation and strong selection drive the nematode to rapidly evolve in adaptation to new environments, which including local pine hosts, vector beetles, commensal microflora and other new environmental factors, during invasion process.

## Introduction

Invasive species cause immense environmental and economic damage worldwide. The pinewood nematode (PWN), *Bursaphelenchus xylophilus* (Steiner et Buhrer) (Nematoda: Aphelenchoididae), is a dangerous invasive species. This migratory endo-parasitic nematode causes a devastating forest disease, pine wilt disease (PWD), and has killed thousands of millions of *Pinus* trees in its introduced regions. Currently, it has become a serious threat to forest ecosystems worldwide^1^ and has been listed as one of the top 10 most important plant-parasitic nematodes^2^.

As they colonize and expand into new ranges, invasive species often exhibit changes in genetic variation, population structure, selection regime and phenotypic traits^3^. The pinewood nematode parasitizes in coniferous trees (primarily *Pinus* spp.), and in nature, it is transmitted by vector beetles (mainly *Monochamus* spp.)^4^. Host resistance^5,6^ and capacity of the vector carrying dauer larvae^7–9^ directly influence the occurrence and prevalence of the disease. Moreover, composition of commensal microflora, mainly endophytic fungi and associated bacteria, also influence the occurrence of the nematode^10–13^. PWN was introduced from North America to Asia and Europe, encountering novel pine hosts, vector beetles and commensal microflora different from those in the native regions, and the nematode has to rapidly adapt to local environments. As a successful invasive species, PWN indeed does well. In mainland China, over 30 years, the occurrence of the nematode spread from the first year only one site^14^ to 316 counties in 16 provinces in 2017 (based on data of the State Forestry Administration of China) (**Fig 1a**), covering a majority of pine tree growing regions (**Fig 1b**). The northernmost sites are in the region of 42 °N latitude (Liaoning province), where the annual mean temperature is less than 7 °C, beyond our previous knowledge that the infected pine trees are asymptomatic when the mean temperature ≤10 °C^15,16^. Most notably, the nematode has obviously overcome the local host’s resistance. It was found that the indigenous species Masson’s pine (*P. massoniana*) has gradually lost its resistance, from a resistant host^5,6,16^ to a susceptible host^17,18^, resulting in the destruction of large tracts of pine forests. In recent years, another indigenous species Chinese red pine (*P. tabuliformis*), one of the most important afforestation tree species in large areas of northern China, has also been infected by PWN^19^. Obviously, the Chinese invasive PWNs exhibit adaptive evolution to fit local hosts.

**Figure 1.**
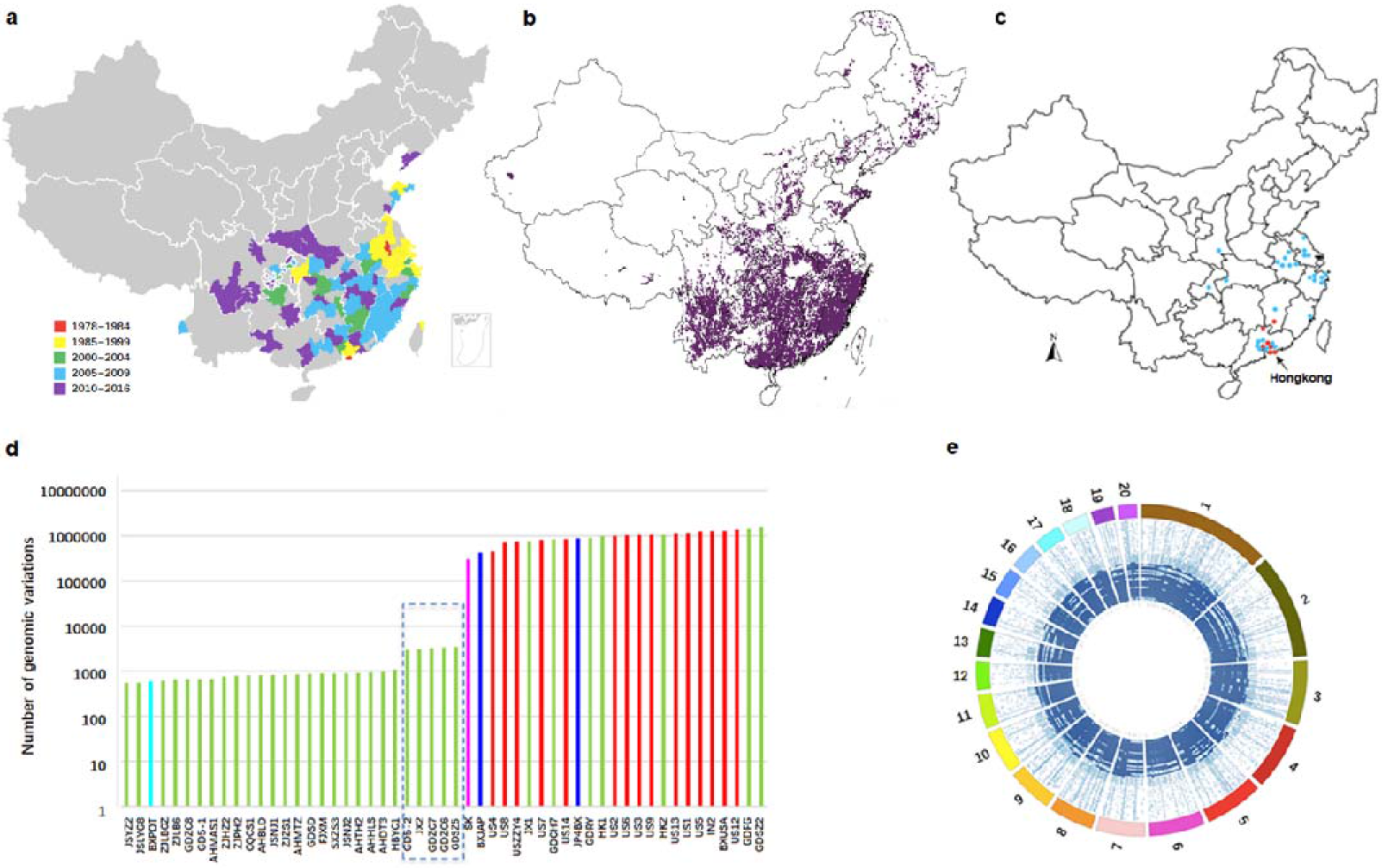
Geographic distributions of the pinewood nematode and pine hosts in China and intraspecific genomic variations of the nematode *B. xylophilus*. a. Distribution map of the pinewood nematode in China. Different colors represent reported years of pine wilt disease epidemic in China, ranging from earlier (red) to more recent (purple) years. b. Distribution map of PWN’s hosts *Pinus* spp. in China. c. Distribution of PWN strains collected in China in this study. Blue dots represent 28 strains comprised the “main group” that are genetically close to each other, while red dots represent other strains showing more genetic diversity. d. Number of genomic variations between each PWN strain and the strain of the reference genome (BxCN). Green, cyan, purple, blue, and red represent strains from China, Portugal, Korea, Japan, and USA, respectively. The strains are arranged based on the number of GVs. The five Chinese PWN strains in the dashed box show low but elevated numbers of GVs. e. Distribution of genomic variations among 55 strains in the *B. xylophilus* genome. Using BxCN as a reference genome, genomic variations of 55 PWN strains mapped on the top 20 scaffolds of the reference genome.

We are interested in the genetic adaptation of PWN to novel environments. Genetic characteristics of populations have profound impacts on their capacity for establishment and range expansions of invasive species^20^. Previous studies on the genetic diversity of PWN showed significant differences among sample sources and detection approaches^21–27^. Most of these studies utilized short genetic markers. Recently, next-generation sequencing has become increasingly accessible and economical, making feasible the genome-wide exploration of genetic diversity and examining the extent of selection in shaping genome-wide levels of genetic differentiation within native and invasive populations. The complete genome sequence of PWN has been published^28^. Taking advantage of this reference genome, a comparative analysis of genome-wide variations of six PWN strains isolated from Japan brought insight into the relationship between high levels of genetic diversity and pathogenic traits^29^. Genetic variation is usually considered to determine the potential for species to evolve in response to new environments^30^; by using whole-genome resequencing of a large cohort of PWN strains isolated from China and other non-native and native regions, we attempt in this study to examine the genome-wide variations of the nematode *B. xylophilus* and to explore genetic underpinnings of successful PWN invasion by comparing genetic differences between the invasive PWNs and the native PWNs.

## Results

### Sequencing and assembly of the reference genome BxCN

To facilitate comparative analysis of PWN strains from different geographic regions, we sequenced and assembled *de novo* the whole genome of one PWN strain isolated from Zhejiang Province, China (named BxCN), and used it as the reference genome. The genome size is 79.2 Mb, with the contig N50 size 43,492 bp and the scaffold N50 size 3.3 Mb (the largest scaffold 8.2 Mb). To ensure high quality of the genome annotation, we applied a three-tier strategy to annotate protein-coding genes in the assembled genome of BxCN: (1) *de novo* annotation, (2) homology-based improvement using the program genBlastG with the published PWN genome protein-coding genes^28^ as queries, and (3) RNA-seq-based gene model validation and further improvement (see Methods). We annotated 17,692 protein-coding genes in BxCN, slightly smaller than the annotated protein-coding genes reported previously from the Japanese strain Ka4 (18,074)^28^ (**Table 1**). The majority of our predicted gene models have been validated and refined using RNA-seq data. Among them, 16,691 genes contain at least one intron, and 11,639 (69.7%) of them have all introns of each gene validated using RNA-seq data. The median gene size (i.e., genomic span) for protein-coding genes is 1,539 bp. The median exon size is 180 bp and median intron 75 bp. We have identified 16,765 genes with orthologous relationship between BxCN and the published *B. xylophilus* genome^28^ (**Fig S1**). At the protein level, the average of ortholog PIDs (protein ID) between the two PWN genomes is 96.62%, while at the nucleotide level, the average of ortholog PIDs is 96.34%. The new genome sequences are deposited in NCBI (accession number: POCF00000000).

**Table 1.**
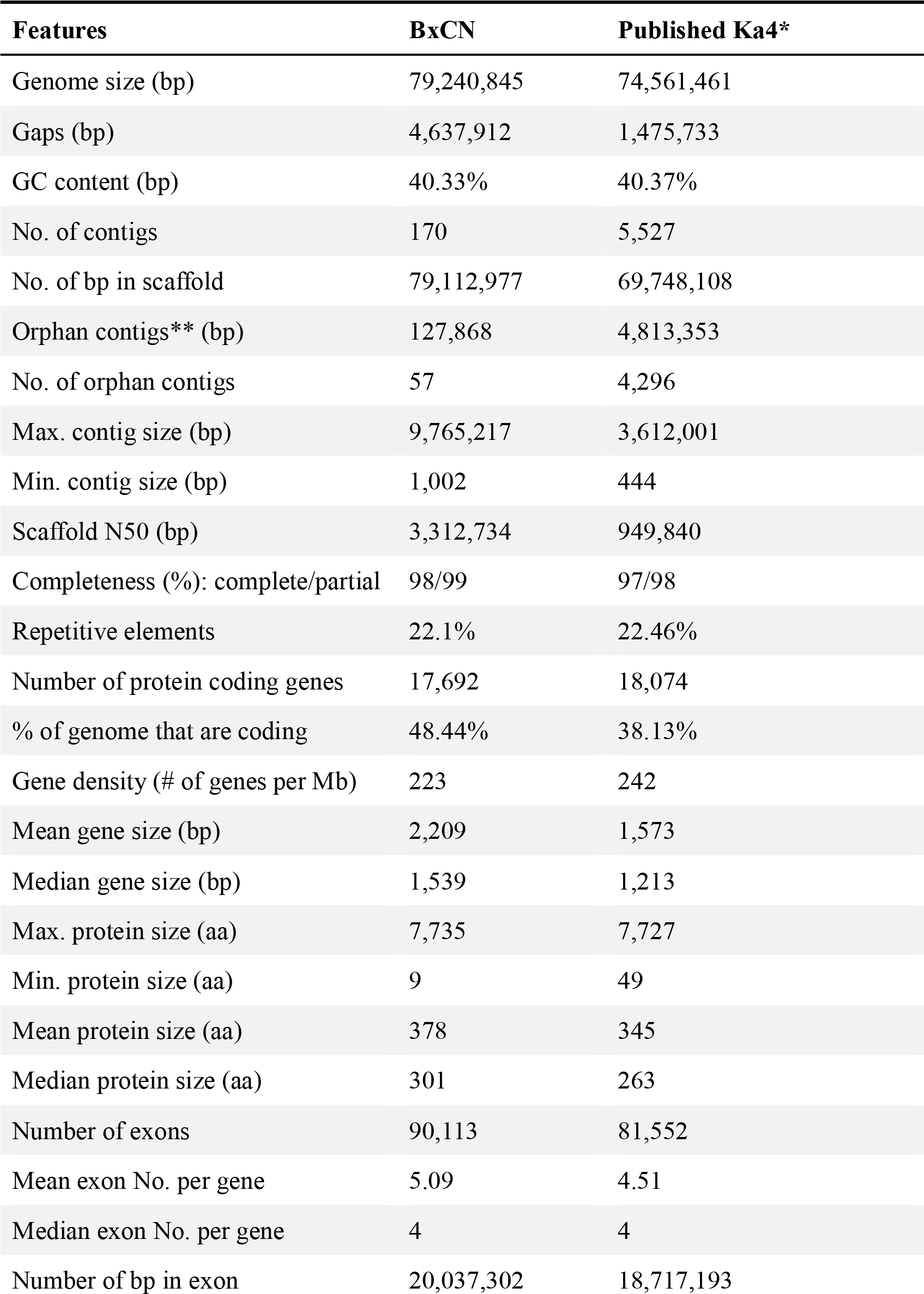
Summary of the newly sequenced genome BxCN compared to the previously published genome of *Bursaphelenchus xylophilus*.

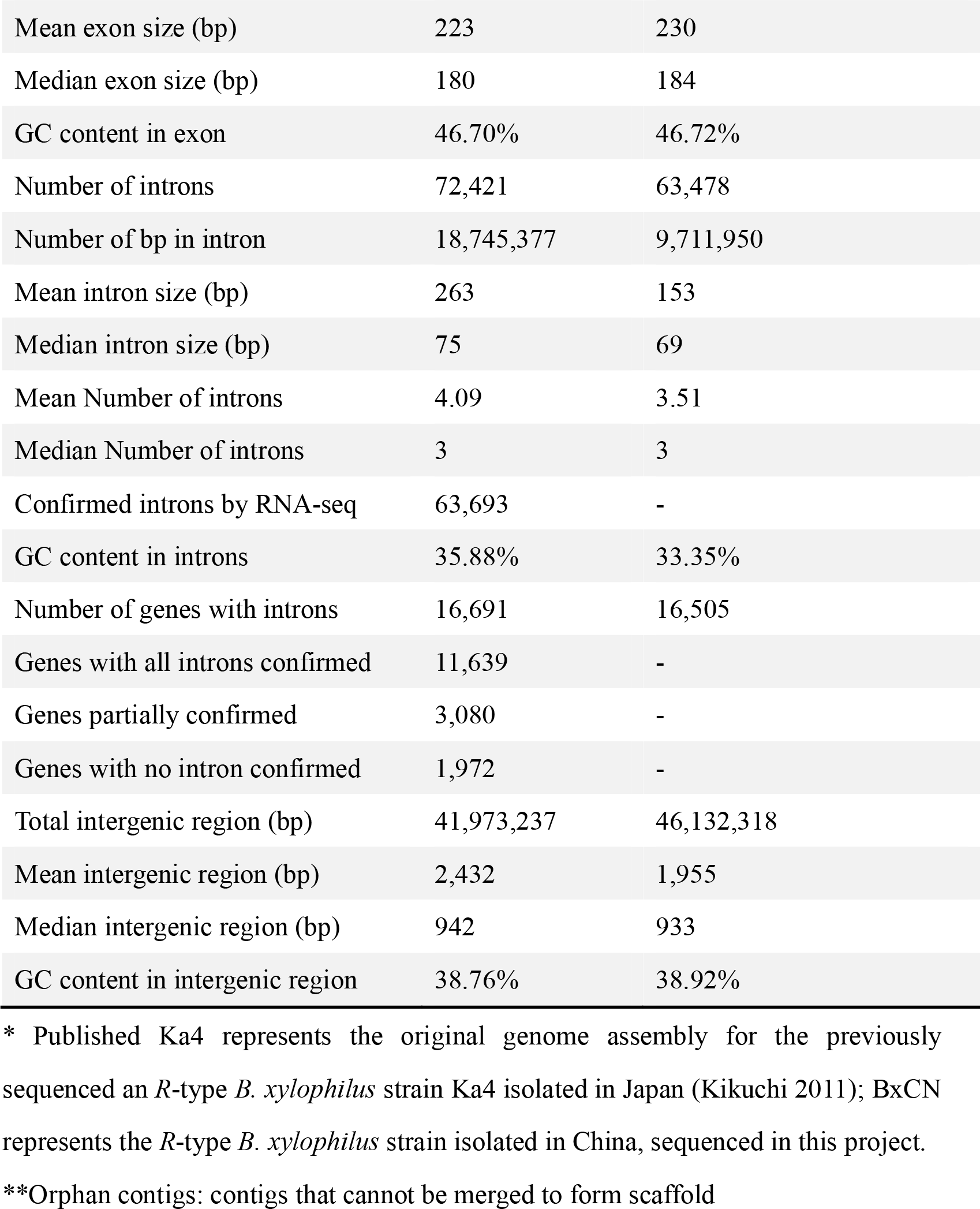

### Whole-genome variation analysis displays significant genetic divergences among PWN strains

In this project, we sequenced the whole genomes of other 54 PWN strains to ~30X coverage (**Table S1**). Among these PWN strains, 35 (including BxCN) were isolated from 10 provinces in China (**Fig 1c**, **Fig S2a**), four from other introduced countries, including two from Japan, one each from South Korea and Portugal, 15 from the native region USA and one from Canada (**Table S1**, **Fig S2b**).

Aligning whole-genome resequencing reads of the other 53 PWN strains (excluding the strain BxCA from Canada, the M-type of *B. xylophilus*, that shared much lower similarity to the R-type strains) against BxCN (mapping rates ranged from 43% to 97%) reveals remarkable differences in the total amounts of genomic variations (GVs, excluded InDels), ranging from a few hundred GVs to over one million GVs (**Fig 1d**), which indicates dramatic genetic divergences among these PWN strains. Most (28 of 35, including BxCN) of the Chinese PWN strains showed fewer pair-wise GVs, suggesting these strains had similar genetic backgrounds. However, the other seven PWN strains isolated from China showed high numbers of GVs (**Fig 1d**), indicating significant genetic divergence between these strains and the reference genome (BxCN). Interestingly, PWN strains with few GVs (blue dots, **Fig 1c**) showed a dramatically different geographical pattern compared to these strains with high GVs in China (red dots, **Fig 1c**), suggesting that PWN strains with different genetic backgrounds might have different abilities in establishment and spread during the invasion process, which might be related to abilities of adaptation to novel environments. We hypothesized that PWN strains isolated from China with few GVs share genetic features that enabled them to spread and adapt to wider geographical regions (**Fig 1c**). Notably, the single Portuguese PWN strain BxPot sequenced in this project also had fewer (595) GVs. In contrast, all PWN strains isolated from the USA and Japan showed much higher numbers of GVs (**Fig 1d**), indicating that they have significant genetic divergences with BxCN. Moreover, we found that the genomic variations are essentially evenly distributed in the *B. xylophilus* genome (**Fig 1e**).

### Phylogenetic relationship inferred from nuclear genomes reveals a dominant invasion group existing in China

To estimate the phylogenetic relationship among PWN strains, we identified genomic regions that have minimal 10× coverage in all strains to ensure high-quality and accurate comparison. We also downloaded the published genome sequences of the Japanese strain Ka4^28^ from GenBank and used for comparison. SNPs located within this “subgenome”, which was 14,981,085 bases in size, were used to construct a phylogenetic relationship of all 55 PWN strains (**Fig 2a**). The phylogenetic tree showed a clade with a strikingly large cluster of PWN strains isolated from China, which also includes the only Portuguese strain, BxPot (**Fig 2a**). The PWN strains within this cluster were exactly the ones that contained few GVs (**Fig 1d**). The other Chinese strains that did not fall into this clustering (7/35) formed separate clusters with other strains, including the genome-published strain Ka4. Because of the extremely close genetic relationship of the PWN strains within the cluster and extremely wide geographical distribution in China (blue dots, **Fig 1c**), and because the group represents the majority of the PWNs isolated from China (28/35), we named it a **d**ominant **i**nvasive **g**roup (DIG). Because this DIG was identified using nuclear DNA sequences, we name it nDIG. The nDIG was composed of 28 Chinese PWN strains and the single Portuguese PWN strain (**Fig 2a**). Interestingly, five PWN strains within the nDIG group formed a minor group (**Fig 2b**), containing strains with slightly higher numbers of GVs (dotted box, **Fig 1d**). These five PWN strains showed low level of pair-wise GVs, suggesting their close genetic relationships. The fact that these 29 PWN strains clustered in a single clade of the phylogenetic tree and the short evolutionary distance among them (**Fig 2a**) suggests that nDIG strains share the most recent common ancestor (MRCA) and that they diverged only recently. Of the nDIG strains, the only PWN strain from outside of China is the one isolated from Portugal (**Fig 2a**, BXPot, blue arrow), suggesting high similarity and a close genetic relationship between the Portuguese strain and nDIG strains isolated from China, supporting the previous hypothesis that PWNs in Portugal might have originated from Asia^31,32^. In contrast, all other PWN strains showed much longer branch lengths in the phylogenetic tree (**Fig 2a**) and had larger numbers of GVs compared to BxCN than the nDIG strains (**Fig 1d, e**). Among the three Japanese PWN strains (**Fig 2a**), JP4BX clustered with the native USA strains, while BXJAP and Ka4 clustered more closely with some non-DIG PWN strains from China, suggesting high genetic differentiation among Japanese strains, consistent with a previous study^29^. The single PWN isolate from South Korea (SK) included in this study was close to but not within the nDIG clad. Despite its relatively close clustering with the nDIG PWN strains isolated from China (**Fig 2a**), the Korea strain showed high genome-wide GVs from BxCN (**Fig 1d**), suggesting that they shared a MRCA and then diverged under different selection regimes. The high sequence similarity of nDIG strains was also revealed in principal component analysis (PCA) (**Fig 2c**).

**Figure 2.**
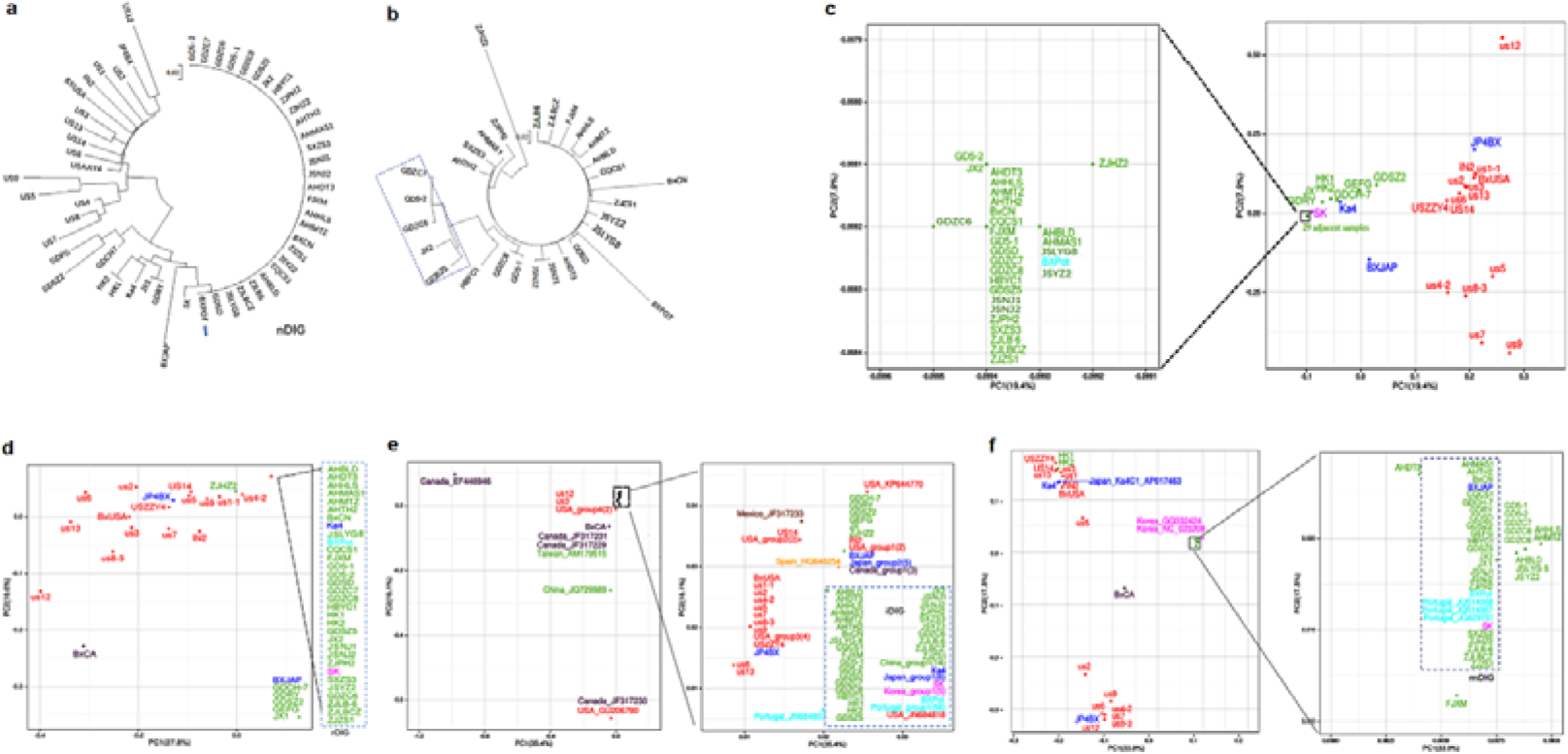
Evolutionary relationship among PWN strains from different geographic regions. a. Phylogenetic tree inferred from nuclear genome sequences of the 55 PWN strains. b. Phylogenetic relationship of the 29 strains comprising the dominant invasive group (DIG). The five PWN strains enclosed in the dotted square formed a clad. c. PCA analysis based on nuclear genome sequences of the 55 PWN strains. d. PCA analysis based on the full-length rDNA sequences. e. PCA analysis based on ITS sequences. f. PCA analysis based on mitochondria DNA sequences.

### Targeted analysis of rDNA sequences hints at the origin of the dominant invasive PWN group

Due to the limitation of samples from outside of China, especially few samples from Japan and other introduced regions that were sequenced and analysed in this project, it is difficult to confirm the origin of nDIG from Japan or USA. To further trace the origin of the nDIG strains isolated from China and define the relationship among PWN strains, we attempted to use rDNA sequences as proxies for whole genome sequences for phylogenetic analysis. We first assembled the full-length rDNA sequences from the genome dataset. PWN full-length rDNA, which consists of IGS, 18S, ITS1, 5.8S, ITS2, and 28S sequences, is 7586 bp in size. We examined how full-length rDNA sequences faithfully represent whole-genome sequences. PCA and phylogenetic analysis of the full-length rDNA sequences in the 56 PWN genomes (including the M-type BxCA from Canada) also display a strikingly similar dominant invasive group (**Fig 2d**, **Fig S3a**). Examination of the full-length rDNA sequences of this DIG (rDIG, for being based on rDNA sequences) revealed that these sequences were 100% identical. As expected, the rDIG and the nDIG were remarkably similar. Slightly different from the nDIG identified using nuclear whole-genome DNA sequences (**Fig 2c**), 32 PWN strains were involved in rDIG (**Fig 2d**). Compared to nDIG, rDIG contained four additional PWN strains, including two Chinese strains isolated from Hong Kong (HK1 and HK2), one Korean strain (SK) and one Japanese genome published strain (Ka4), indicating that the rDNA sequences of these four PWN strains provides relatively lower resolution in distinguishing PWN strains than the whole-genome sequences. Interestingly, a Chinese strain ZJHZ2, which is a member of the nDIG (**Fig 2c**), was absent from the rDIG (**Fig 2d**). Examination of the rDNA sequence of ZJHZ2 and other PWN strains in rDIG showed that four SNPs in ZJHZ2 were different from other strains (**Fig S4**). Because of the extremely high conservation of rDNA sequences in evolution, the differences in rDNA sequences between ZJHZ2 and rDIG strains suggest that rDNA sequences in ZJHZ2 might have a different origin from its full-length nuclear genome. In other words, the ZJHZ2 strain might be the progeny of an ancestral nDIG strain and an ancestral non-nDIG strain, implying introgression existed in PWNs. Our result suggested that despite its value in phylogenetic analysis, rDNA sequence may not always represent the whole-genome sequences. Taken together, full-length rDNA can basically represent the whole-genome sequences reasonably well, with a few important exceptions.

Then, we downloaded the published PWN rDNA sequences from NCBI. We focused on the ITS sequence components of the rDNA sequences, as most previous projects only sequenced the ITS region (including ITS1, 5.8S, and ITS2 regions) of the rDNA sequence, which is 784 bp in size, representing a small but important portion of the full-length rDNA sequence. We catalogued all ITS sequenced obtained in this project, together with all previously published ITS sequences from PWN strains (**Table S2**) and used them to examine the evolutionary relationship among the PWN strains. Altogether, 167 full-length ITS sequences were obtained, with 55 generated in this project and 112 retrieved from previously published results. Of these, 50, 14, 6, 58, 8, and 28 sequences were from strains isolated from Mainland China, Japan, Korea, Portugal, Canada, and the USA, respectively. The other three ITS sequences were from strains isolated from Mexico (1), Spain (1), and Taiwan, China (1). As expected, PCA and phylogenetic analysis showed one large group, named iDIG (for being based on the ITS sequences), which included all 32 PWN strains in the rDIG group and other 81 additional PWN samples (**Fig 2e**, **Fig S5**). Notably, among those additional PWN samples, 56 were from Portugal, 14 from China, 6 from Japan, 5 from South Korea, and one from the USA (JN684818)^32^. All these ITS sequences share 100% identical sequence. The inclusion of one PWN strain isolated from the USA indicates that the ancestor of iDIG existed in the native USA.

### Analysis of mitochondrial genomes indicates intraspecific genetic admixture existing in DIG ancestor strains

Mitochondria genomes are inherited from the maternal ancestor. To explore the maternal inheritance of PWN strains, we analysed 62 mitochondrial genomes, among which 55 were constructed in this project and six were obtained from GenBank, which includes strains from Portugal (3), South Korea (2) and Japan (1). Similar to previous analyses, PCA and phylogenetic analyses of these 62 mitochondria genomes revealed a dominant invasive group (mDIG, as identified through analysing mitochondrial genome sequences) (**Fig 2f**, **Fig S3b**). The mDIG contained 27 mitochondria genome sequences from PWN strains isolated from China (21), Portugal (4), Japan (1), and Korea (1), suggesting they share maternal MRCA. Notably, the 21 Chinese strains in mDIG were not identical to those in nDIG. Except from two strains from Hong Kong (HK1, HK2), the other five non-nDIG strains were involved in mDIG (JX1, GDRY, GDSZ2, GEFG, GDCH7), but 12 nDIG members (FJXM, AHDT3, AHHLS, AHMTZ, AHBLD, JSLYG, JSYZ2, GD5-1, GD5-2, GDZC6, GDZC7, GDZC8) were very close but differentiated from mDIG. Interestingly, ZJHZ2 was identified as a member of mDIG (**Fig 2f**) and was also identified as a member of nDIG (**Fig 2c**), but not identified as a member of rDIG (**Fig 2d**, **Fig 2e**), suggesting that the majority of genetic material of ZJHZ2 is inherited from DIG ancestors. Additionally, the Korean strain SK and the Japanese strain BxJAP are not in nDIG but are in mDIG. Based on these important differences between the mDIG and nDIG, we suggest that intraspecific genetic admixture occurred in DIG ancestor strains, resulting in paternal and maternal inheritance that are not always identical in DIG PWNs.

### Genetic drifts and selection are primary evolutionary factors shaping genetic diversity patterns of PWN populations

To ensure high quality, we focused the analysis on nucleotides that satisfied the strict criteria (reads supporting ≥90%, coverage ≥10). A total of 169,388 credible SNPs were determined. We noticed that most of these GV sites (125,199 SNPs, 74% of total) were polymorphic among the 15 USA strains, indicating that the native USA population has rather abundant genetic variations. Approximately 31% sites (51,785 SNPs) were polymorphic among the three Japanese strains, which was more than those (32,801 SNPs, 19% of total) among the 35 Chinese strains. Notably, few sites (478 SNPs, <0.3%) were polymorphic among the 29 nDIG strains, and the overwhelming majority of those were GV sites with fixation alleles (100% frequency) in nDIG PWNs.

We paid more attention to variation sites in which low frequency alleles (<50%) in the native USA strains were fixed in DIG strains, and a total of 25,992 SNPs (15.3% of total) screened out (**Fig 3a**). We found that the majority of these SNPs belonged to standing variations (**Fig 3b**), of which approximately 40% of DIG-fixation alleles were pre-existing in the 15 USA strains (10,530 SNPs), 50% in the three Japanese strains (12,949 SNPs) and 2.5% in the seven Chinese non-DIG strains (662 SNPs). We suggest that whether these fixation alleles originated from the native populations or invasive populations due to founder effects and genetic drifts, in addition to selection action in new environments, these alleles have been fixed in nDIG strains. The other DIG-fixation alleles (1851 SNPs, approximately 7% of total) were specific (**Fig 3b**), and we have not yet determined their origins. Of them, 32 alleles were fixed in all invasive strains (including the Chinese, Portuguese, South Korean and Japanese strains), 508 alleles in invasive strains excluding the Japanese PWNs (i.e., the Chinese, Portuguese and South Korean strains, named CH-Port-SK group), 1144 alleles in nDIG and the South Korean strains (named DIG-SK group), and 167 alleles only in nDIG strains. We hypothesized that these variations might originate from new mutations during the invasion process. Alternatively, they might be pre-existing variations, but not be detected in this study because of sampling bias. Whether they belong to novel mutations, standing variations or both, under selection action, these genes have been fixed in different groups.

**Figure 3.**
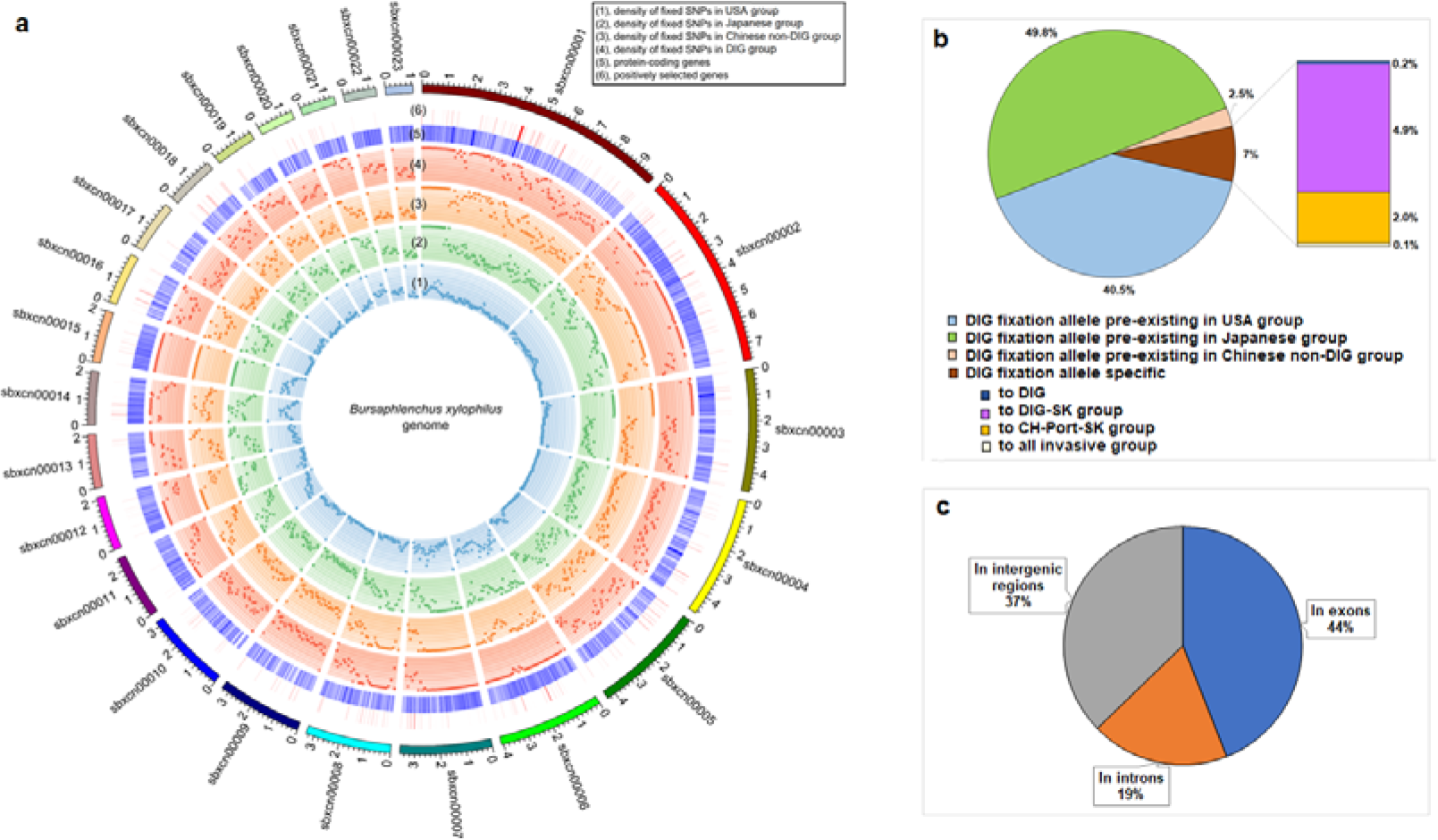
Distribution of genomic variations with low frequency alleles (<50%) in the native USA group but fixation in the DIG (25,992 SNPs) a. Density of SNPs fixed in different PWN groups mapping on the top 23 scaffolds of the reference genome. Each dot represents density of fixed SNPs within a 100Kb genomic region. That was calculated with a formula: the number of fixation SNPs /total SNPs. b. Variation types of DIG-fixation SNPs, showing majority of them belong to standing variation. c. Proportion of DIG-fixation SNPs locating in coding, non-coding and intergenic regions.

We checked the location of these 25,992 SNPs in the reference genome (BxCN) and found they are unevenly distributed in the genome (**Fig 3a**), different from the total SNPs distribution in the genome. Fixation alleles specific to nDIG strains (167 SNPs) were mainly located in scaffolds sBxCN00003 (54.5%) and sBxCN00007 (18.6%); those specific to DIG-SK strains (1144 SNPs) mainly in scaffolds sBxCN00015 (41.3%) and sBxCN00003 (37.6%), as were those specific to all invasive strains (32 SNPs), while those specific to CH-Port-SK strains (508 SNPs) were mainly in scaffolds sBxCN00007 (71.6%), sBxCN00011 (12.6%) and sBxCN00001 (10.4%). Interestingly, the above three scaffolds also harbour nearly 65% of SNPs with fixation alleles pre-existing in the Chinese non-DIG strains (662 SNPs). Those with fixation alleles pre-existing in the Japanese strains (12,949 SNPs) were relatively rich in the five scaffolds (sBxCN00002, sBxCN00003, sBxCN00015, sBxCN00001 and sBxCN00007, in order), which harbour more than 58% of those GVs. Comparatively, distribution of SNPs with fixation alleles pre-existing in the USA strains (10,530 SNPs) were more even in the genome, slightly rich in the five scaffolds (sBxCN00002, sBxCN00001, sBxCN00008, sBxCN00006 and sBxCN00005, in order), which harboured nearly 50% of those GVs. The result indicates selection acting on different genomic regions, perhaps accompanied by selective sweeps, which might be related to adaption of the nematode to local environments.

### Functional annotation of GV-harboured genes reveals genetic adaptation of the nematode *B. xylophilus*

Functional annotation of GV-harboured genes may help us to understanding genetic mechanisms of successful PWN invasion. Based on the locations of these 25,992 SNPs in the reference genome (BxCN), relative genes were identified. A total of 16,312 GVs were in coding regions, harboured within 6580 genes (including 11443 GVs in exons and 4869 GVs in introns), and the other 9681 GVs were in the intergenic (IG) regions (**Fig. 3c**). Among those genes, nearly 70% (4601) had a Pfam annotation (**Table S3**). Protein kinase domain (PF00069) was most abundant, followed by MFS (major facilitator superfamily) (PF07690), 7TM GPCR rhodopsin family (PF00001), protein tyrosine kinase (PF07714), and neurotransmitter-gated ion-channel ligand binding domain (PF02931), in order. The top 35 Pfams are shown in **Fig 4a**. We noticed that chemoreceptors (16 families 225 genes); proteases (38 families 213 genes); transport proteins (e.g., MFS 58 genes, ABC transporters 7 families 39 genes); carbohydrate-active enzymes, such as glycosyl hydrolases (12 families 23 genes) and glycosyl transferases (13 families 32 genes); detoxification enzymes, such as cytochrome P450 (CYPs, 26 genes), short chain dehydrogenases (SDRs, 26 genes), UDP-glucuronosyl/UDP-glucosyl transferases (UGTs, 17 genes) and glutathione S-transferases (GSTs, 3 families 11 genes); and nematode cuticle collagen N-terminal domain (22 genes) were abundant. These genes have important biological functions. Gene Ontology (GO) annotation shows that 4024 genes matched to GO terms. The most abundant term was protein binding (GO:0005515) in the molecular function category. Extracellular vesicular exosome (GO:0070062), cytosol (GO:0005829) and membrane (GO:0016020) were the top three terms in the cellular component category. In the biological process category, single-organism cellular process (GO:0044763), regulation of biological quality (GO:0065008), organ development (GO:0048513), signal transduction (GO:0007165) and positive regulation of cellular process (GO:0048522) were the top five terms. Notably, many genes were assigned to terms of response to stimuli and stresses. Among the top 35 terms in the biological process category, eight terms were related to response to various stimuli and stresses (**Fig 4b**), including response to stress (GO:0006950), response to stimulus (GO:0050896), response to oxygen-containing compound (GO:1901700), response to drug (GO:0009605), response to external stimulus (GO:0042493), response to chemical stimulus (GO:0042221), response to organic substance (GO:0010033) and response to abiotic stimulus (GO:0009628). More than one-third of the annotated genes (1393) were assigned to GO terms related to response to different stresses, stimuli and substances, internal or external. Additionally, many genes were involved in the positive or negative regulation of different biological processes. KOG annotation shows that, except for R (general function prediction only), T (Signal transduction mechanisms) was the most abundant cluster (**Fig 4c**). KEGG annotation showed that these genes were mainly involved in metabolisms of carbohydrates, lipids, amino acids, xenobiotics biodegradation, glycan biosynthesis, and signal transduction (**Fig 4d**).

**Fig. 4.**
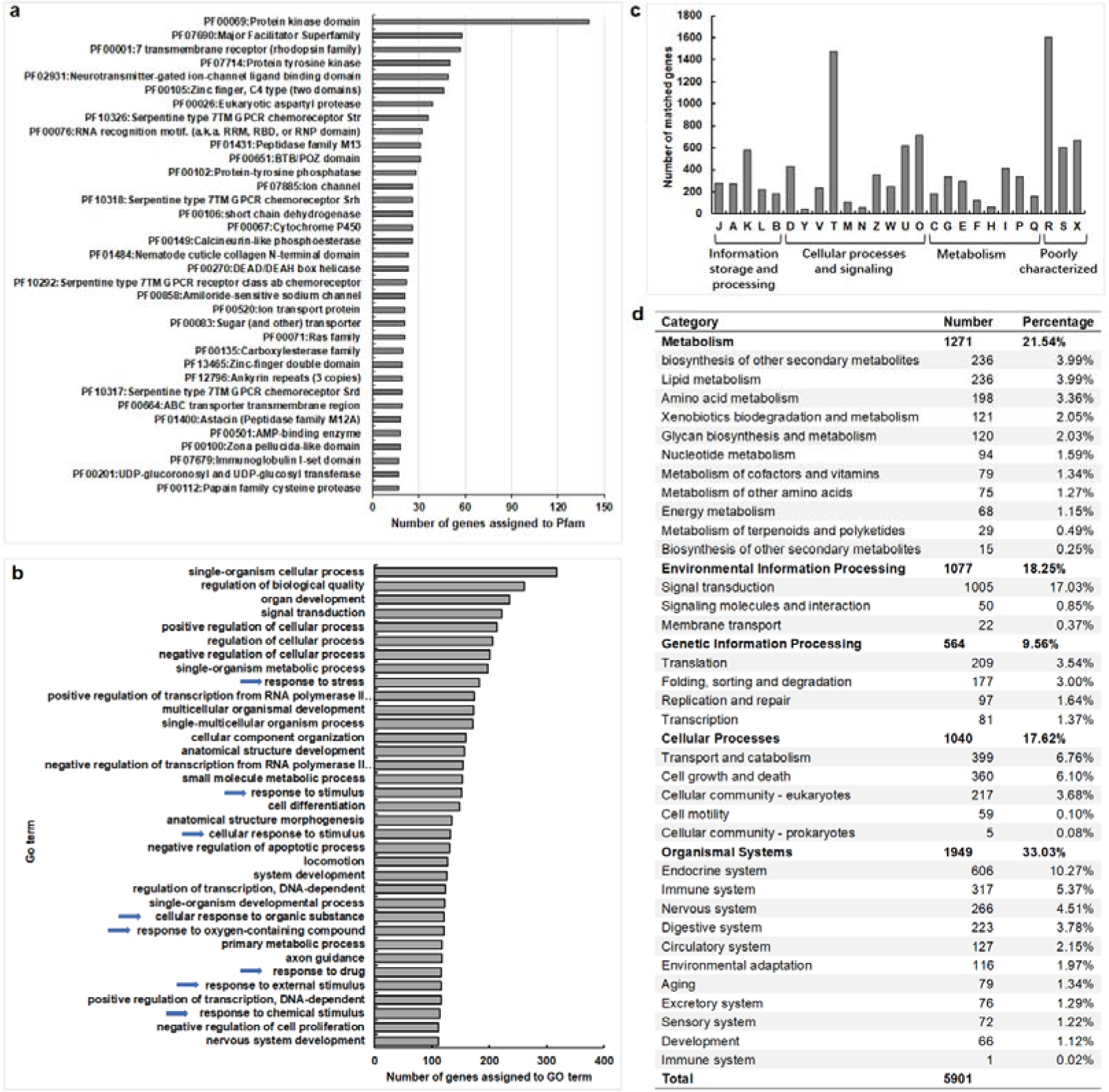
Functional annotations of protein-coding genes harbored DIG-fixation SNPs. a. The top 35 Pfams. b. The top 35 GO terms in the biological process category. c. KOG annotation. d. KEGG categories of genes assigned.

We paid more attention to those genes that harboured GVs with specific alleles (**Fig 3b**), which are perhaps more likely to be related to local ecological adaptation and invasiveness of the nematode. Among those 1851 SNPs, 1201 were harboured within 626 genes (including 862 GVs locating in exons and 339 GVs in introns).

Among them, 18 genes had fixation alleles specific to all invasive strains, 178 genes specific to invasive CH-Port-SK strains, 370 genes specific to nDIG-SK strains, and 60 genes specific to nDIG strains. Most of these genes had a functional annotation. Among them, the most abundant Pfams were protein kinase domain (PF00069), zinc finger C4 type (PF00105), short chain dehydrogenase (PF00106), and major facilitator superfamily (PF07690), in order. Based on their functional annotation, we divided genes into the following groups, which encode proteins putatively related to adaptation (**Table S4**). **Group 1**, G-protein-coupled receptors (GPCRs). Eight serpentine type 7TM GPCR chemoreceptors were involved, including two from family Srt, and one from each family Str, Srv, Srh, Srbc, Srab and Srsx. Because chemicals produced by pine hosts and vector beetles can regulate nematode behaviours in the life cycle of *B. xylophilus*^33,34^, these GPCR chemoreceptors may play important roles in chemical communication between PWN and local hosts, as well as local vector beetles. Moreover, four rhodopsin family GPCRs were found. Rhodopsin GPCRs were reported to play important roles in regulation of food intake in nematodes^35^. **Group 2**, proteases. A total of 19 peptidases were involved, including two cysteine proteases, two eukaryotic aspartyl proteases, three serine carboxypeptidases, one tripeptidyl-peptidase, and one rhomboid protease, as well as ten metallo peptidases (including three family M13 and one from each family M28, M20, M1, M24, M12, ubiquitin protease and zinc carboxypeptidase). Proteases mainly play roles in digestion and acquisition of nourishment, and some other functions. **Group 3**, detoxification enzymes. This group included three SDRs, three UGTs and one GST, which may be involved in cytochrome P450 xenobiotics metabolism; three of each aldehyde dehydrogenases and enoyl-CoA hydratases/isomerases, two of each FMOs, dihydropyrimidinases and acetyl-CoA acetyltransferases, and one of each thymidine kinase, IMP dehydrogenase and GMP synthase, which are putatively involved in drug and other xenobiotics metabolisms; eight ABC transporters and five MFS proteins, which may also play roles in detoxification by extruding toxins and drugs out of the cell. **Group 4**, proteins involved in signal transduction. This group included eight zinc finger proteins (including six C4 type, two C2H2 type and one C3H4 type), four Ras family proteins, 12 protein kinases and four protein tyrosine kinases, two low-density lipoprotein receptors, and one of each glutamate receptor, Rap/ran-GAP family protein and paracaspase, which may play important roles in signal transduction. **Group 5**, proteins involved in response to stresses and stimuli. This group included one catalase, three acid phosphatases and one serine/threonine-protein phosphatase, three haloacid dehalogenase-like hydrolases, and one of each heat shock protein Hsp20 and DnaJ domain protein, might be related to response to stresses and stimuli. **Group 6**, other important proteins. These proteins included four gamma interferon inducible lysosomal thiol reductases (GILT), which might be involved in immune system; two regulators of chromosome condensation (RCC1) that may play an important role in the regulation of gene expression; two chitin synthases, one alpha-IV type collagen and three nematode cuticle collagens that may be related to the formation of the nematode cuticle, which is a physiologically active structure involved in a variety of functions, such as absorption, excretion, transport, locomotion, and as a protective barrier. All of the above proteins may play important roles in the nematode adaptation to local environments.

### dN/dS test shows positive selection acting on functional important genes

We calculated the ratio of non-synonymous (dN) to synonymous (dS) substitutions of all those 6580 genes, and 221 genes were predicted to be under positive selection (dN/dS > 1, P-value < 0.05). Of these, 151 genes had a functional annotation (**Fig 3a**, **Table S5**). Important functional genes comprised eight GPCRs (including three family Srab, three family Str, one family Srw and one G-protein coupled receptor frpr-1 homologue); twelve proteases (including six family M13, and one of each family M1, C13, S08, S28, rhomboid protease and eukaryotic aspartyl protease); six protein kinases and a tyrosine kinase; detoxification enzymes (including two cytochrome P450, one of each UGT, SDR, GST and aldehyde dehydrogenase, two FMO-like proteins, one ABC transporter and two MFS proteins); and other important proteins, such as three zinc-finger proteins, three RNA recognition motifs, two pre-mRNA-processing factors, two lipases, three immunoglobulin I-set domains, three glycosyltransferases, two carboxylesterases, and one transthyretin-like family protein.

Among these positively selected genes, 17 genes harboured GVs with specific alleles (**Table S5**). Of them, three are specific to nDIG strains, including one of each eukaryotic aspartyl protease, FMO-like protein and initiation factor. Ten were specific to DIG-SK strains, including one of each rhomboid protease, aldehyde dehydrogenase, ABC transporter, collagen alpha-1, CaiB/baiF CoA-transferase, GPI ethanolamine phosphate transferase, neurotransmitter-gated ion-channel ligand binding domain, and three functional unknown proteins. Three were specific to CH-Port-SK strains, including one serine/threonine-protein kinase, one ATP-sulphurylase and a functional unknown protein. One unknown function protein was specific to all invasive strains. All these positive selection genes may play important roles in PWN adaption to novel environments.

## Discussion

Determining the source populations and routes of invasion is important for control of invasive species. Much effort was made to map and understand the origin and migration paths of different invasive PWN populations by using various genotyping methods in previous studies^22,25,31,32,36,37^, but the results are still unclear or controversial. In this project, through whole genome sequencing and comparative analysis of a large cohort of PWN strains isolated from different regions, we clearly defined two groups of PWN strains within China, the main group (DIG) and minor group (non-DIG), with striking geographical distribution (**Fig 1c**). The distribution of PWN strains of the DIG correlated with the rapid expansion of forests with pine wilt disease (**Fig 1a**). Analysis of data collected in this study and historical data obtained in public databases (**Fig 1a**) indicated a rapid invasion and migration of PWN in China, which is consistent with the high similarity of genomic sequences of PWN strains in the DIG (**Fig 1c**), suggesting that a dominant invasive population has formed in China. Our comprehensive analysis revealed two clear migration paths of PWN in China (**Fig 5a**): one followed by the DIG strains (blue line) and another one followed by the non-DIG strains (red line). Notably, the dispersal path of DIG in China in this study is similar to one path proposed in a previous study, which suggested that PWN was spread from Guangdong to Jiangsu, and then from Jiangsu to other regions, such as Hubei, Guizhong and Congqing, and suggested that Jiangsu has become a new spreading centre^25^. Perhaps because of the ‘bridgehead effect’ of Jiangsu as a spreading centre, ten years later, the spread population has become a dominant invasive population, namely, the current DIG. To our surprise, Portuguese PWN strains also gathered into the DIG clade, indicating the dominant invasive population that has introduced to Portugal. Moreover, the South Korean strain is closer to DIG than to other non-DIG strains, indicating that the South Korean PWNs and the Chinese DIG strains have close relationship. Our result is similar to that obtained from analysis of effector transcript SNPs in a previous study, which showed a closer proximity of the Portuguese isolates to the Korean and Chinese isolates than to the Japanese or American isolates^32^. Although based on our current genome dataset it is difficult to determine the source population of DIG, targeted analysis of rDNA sequences of PWN strains showed it originated from Japanese but could be traced to the native US population. Based on comprehensive consideration of phylogenetic relationships inferred from nuclear genome, mitochondrial genome, rDNA and ITS sequences, the global spread routes of DIG are proposed, i.e., the invasive PWNs originated from United States and were first introduced to Japan, then to China and South Korea, and finally to Portugal (**Fig 5b**). This finding is identical to the initial knowledge of the PWN migration trajectory around the globe obtained from reported occurrences of PWD in different countries: in the 1900s, it entered into Japan^38^; in the 1970s, it entered into Hong Kong of China^39^; in the 1980s, it entered into mainland China^14^, Taiwan of China^40^ and South Korea^41^, in order; and in the 1990s, it entered into Portugal^42^ and its adjoining regions^43^.

**Fig 5.**
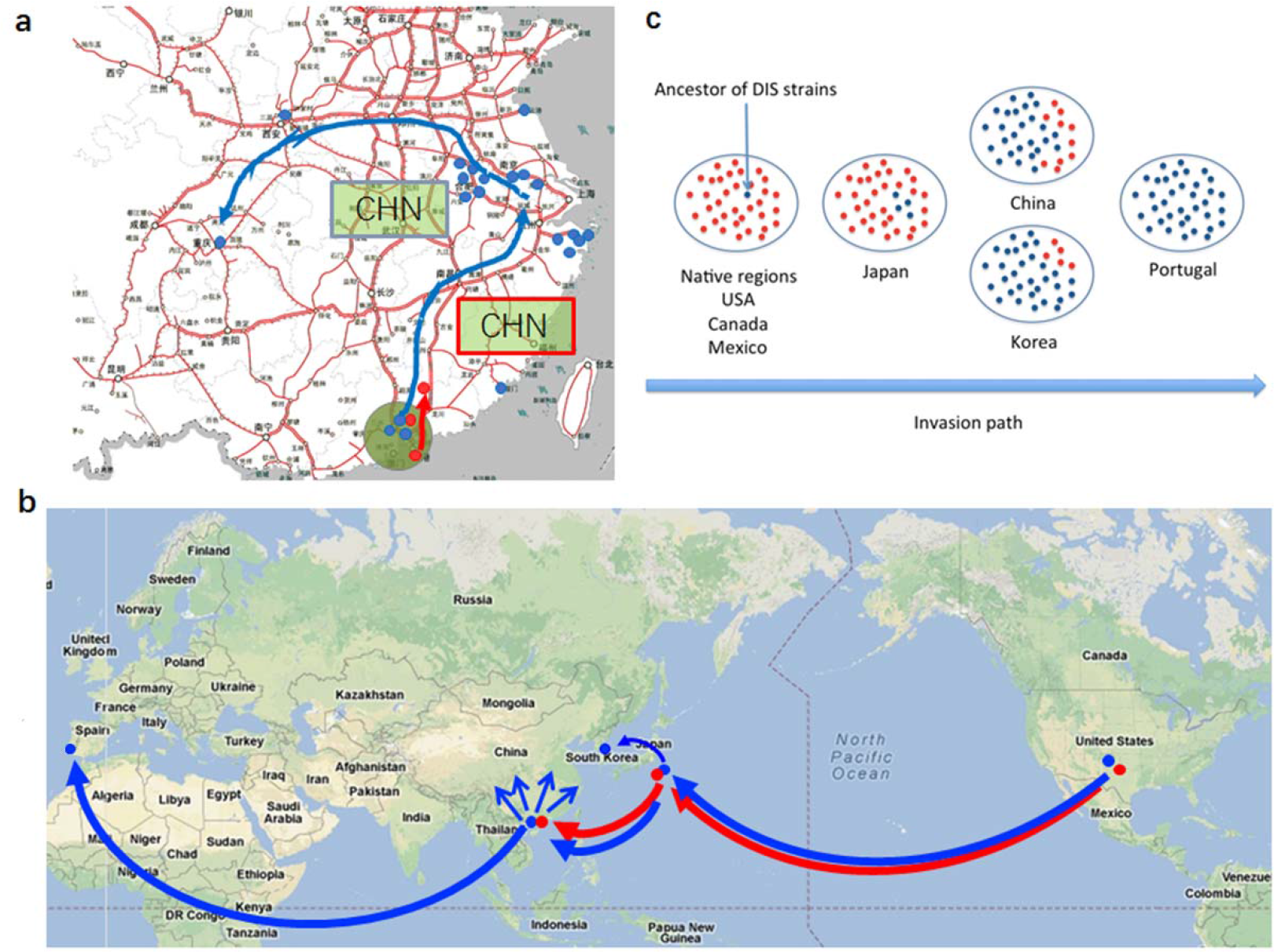
PWN migration paths in China and the globe, with a progressive loss of genetic diversity along the invasion path. (a) Within Mainland China, PWN entered China through Hong Kong and migrated to other provinces via two routes. The “major group” (DIG) migrated to other province along the blue line (blue dots), while the “minor group” (non-DIG) migrated along the red line (red dots). (b) Global migration paths of PWN, from native countries, through non-native country Japan, to South Korea, China, to European countries Portugal and Spain. (c) Losses of genetic diversity along the PWN global migration paths. Red dots indicate non-DIG strains that are genetically diverse, while blue dots indicate DIG strains that are genetically homogeneous. Along the migration paths, percentages of PWN strains included in the DIG is an increase, indicating genetic diversity is a progressive decrease.

Along with PWN migration paths, a progressive loss of genetic diversity was observed. In this study, among the 54 PWN strains (excluding the M-type strain BxCA from Canada), 4,572,007 GVs (including indels) were detected (read supporting ≥60%, reads coverage (i.e., depth) ≥6), approximately 5.8% of the whole genome of *B. xylophilus* (BxCN 79,233,952 bp). Abundant genetic variations in the PWN genome indicate that the nematode has potential to adapt to broad environments. Moreover, we found that the majority of SNPs exist among the native USA strains.

Among those 169,388 credible SNPs (read supporting ≥90%, reads coverage ≥10), nearly 74% GVs are polymorphic among the 15 USA strains, indicating that the genetic diversities are abundant in the native USA PWN population, identical with the results in previous studies^21,22^. High genetic diversity also exists in the Japanese PWN population. Among those GV sites, 31% were polymorphic in the three Japanese strains, obviously higher than that in Chinese PWNs, which was only 19% in the 35 strains. Rich genetic diversity in Japanese populations was also detected in a previous study, which reported that approximately 4.1% of the genome positions were variable as SNPs or indels in six Japanese PWN strains^29^. Notably, the genetic diversity in Chinese PWN strains was mainly in non-DIG strains, while genetic diversity in DIG strains was extremely low, which is the same as the finding in Portuguese PWNs^26,27^. The overwhelming majority of SNPs are fixed in DIG, indicating obvious genetic bottleneck and strong selection, even selective sweep acting on DIG strains. Assuming that the sampling of PWN strains in each country is representative and sampling accurately reflects the actual genotypes of each country, based on ITS sequence dataset, we found that 86% (43/50) of PWN strains isolated from China were included in the iDIG. In comparison, 98.3% (57/58) of Portugal, 100.0% (6/6) of Korea, 50% (7/14) of Japan, and 3.6% (1/28) of USA were members in the iDIG (**Fig 2e**, **Fig S5b**). Thus, from the regions with most recent report of PWN invasion (including Portugal, Korea, China and Japan) to the native USA, there is a decreasing trend in percentages of PWN strains included in the iDIG. The above results indicate that along with the migratory path, i.e., from the USA, to Japan, China, Korea and Portugal, in order, the genetic diversity of PWN clearly displays a progressive decrease (**Fig 5c**).

Genetic diversity determines evolutionary potential and adaptability to new environments of an invasive species. Populations adapt to novel environments in two distinct ways: selection on pre-existing genetic variations (standing variations) and selection on new mutations^44^. We found that a vast majority of fixation alleles in DIG strains were pre-existing in the native USA strains, the Japanese strains or the Chinese non-DIG strains (**Fig 3b**), indicating that these GVs are standing variations. Standing variations have an important role in facilitating rapid adaptation to novel environments because they are immediately available and their functions have been pre-tested^44^. In addition, a small part of fixation alleles (7%) are source undetermined (**Fig 3b**). Although genetic drift could not be excluded, we believe that some of these should originate from novel mutations. Selection acts on both standing variations and novel mutations, shaping patterns of PWN genetic diversity. We also noticed that those GVs with specific alleles were intensively located in a few scaffolds (**Fig 3a**), suggesting strong selection or selective sweep acting on certain genomic fragments. Moreover, 221 GV-harboured genes are predicted under positive selection. Functional annotation of all GV-harboured genes showed that chemoreceptors, proteases, detoxification enzymes, proteins involved in signal transduction and response to stimuli and stresses are abundant, which are important for the nematode’s adaptation to local hosts, vector beetles and other environmental factors in novel ecosystems. However, some pathogenic related genes, such as genes encoding cellulases, pectate lyases, and expansins, which are important for degradation of plant cell walls and related to pathogenicity, are very few, indicating that invasiveness of PWN is related to adaptability to new environments rather than to pathogenicity. Our results shed light on genetic mechanisms of PWN invasiveness.

In conclusion, abundant genetic diversity in native populations provides material of rapid evolution, genetic drift and strong selection as the primarily evolutionary factors that, in addition to mutation, have driven the nematode to rapidly evolve to adaptation to new environments during invasion process. This evolution is the nature of invasiveness of the nematode *B. xylophilus*.

## Materials and Methods

### PWN strains collection

We isolated 35 PWN strains from infested pine trees in 10 provinces in China (**Fig S2a**). We also isolated 20 PWN strains in pine trees from other countries directly collected or intercepted by the quarantine departments (**Fig S2b**). More details about the PWN strains are described in **Table S1**. After isolation from infested wood and identification by morphology under a microscope, nematodes of each strain were sterilized with 3% H_2_O_2_ for 10 min and 0.1% streptomycin sulphate at 4 °C overnight, and then cultured on the fungal mat of *Botrytis cinerea* grown on potato-dextrose agar (PDA) plates containing 0.1% streptomycin sulphate at 25°C. After molecular identification by ITS-PCR-RFLP^45^, cultured nematodes were starved at room temperature for 24 hours and prepared for DNA extraction. All strains were the ‘R-form’ of *B. xylophilus*, excepting the strain from Canada (BxCA), which was the ‘M-form’. For the strain used for deep sequencing and taken as a reference genome (BxCN), sister-brother mating was performed over ten generations, and an inbred line was established for DNA extraction.

### Genomic DNA isolation, library construction, sequencing and assembly

DNA extraction used a regular phenol/chloroform procedure. Briefly, the procedures are as follows: Nematodes were suspended in five volumes of extraction buffer (100 mM Tris-HCl pH = 8.0, 200 mM NaCl, 50 mM, ethylene diamineteraacetice acide (EDTA), 1% sodium dodecyl sulfate (SDS), 1% β-Mercaptoethanol, and 2 mg/ml proteinase K), and incubated at 65 ◦C for 1 h with occasional mixing. The DNA was extracted twice with an equal volume of phenol: chloroform: isoamyl alcohol (25:24:1), and then twice with an equal volume of chloroform: isoamyl alcohol (24:1). The DNA was recovered by ethanol precipitation.

Genomic DNA from each strain was randomly fragmented, and DNA fragments of specific length were gel purified and used for library construction. Libraries with insert sizes for each nematode strain are shown in **Table S1.**For effective genome assembly of the reference genome, multiple libraries with different insert-size (350 bp, 700 bp, 2 kb, 5 kb and 10 kb) were constructed for BxCN. Adapter ligation and DNA cluster preparation were performed and then subjected to Genome Analyzer for pair-end sequencing using Illumina sequencing platforms (GA II or HiSeq 2000 sequencing systems) with standard procedures at the Beijing Genomics Institute (BGI), Shenzhen, China.

The sequence data from each library were filtered to remove low quality sequences, base-calling duplicates and adaptor contamination. For the reference genome BxCN, genome sequences were assembled using SOAPdenovo with default parameters^46,47^. We first assembled the reads into contigs from the short insert-size libraries (≤500 bp) using *k*-mer overlap information (*de Bruijn* graph *k*-mer). The resulting contigs were checked for uniqueness by determining an unambiguous path in the *de Bruijn* graph. Then, we mapped all available paired-end reads to these contigs for connecting adjacent contigs. A hierarchical assembly method was used to construct scaffolds step by step, by adding data from each library separately in the order of insert size from smallest to largest. Gaps were filled in by following the methods described previously^47^. Then, through a reference-based procedure, we attempted to improve BxCN genome assembly. We used the published Japanese PWN genome assembly as the reference^28^ (accession numbers: CADV01000001-CADV01010432), aligned BxCN scaffolds/contigs using mugsy, a genome alignment program^48^. To reduce random alignment events, we used–c (minimum alignment block) value of 500 bp and–d (maximum InDels in alignment block) value of 10,000 bp. With the alignment, we first defined collinear block for every reference/query scaffold/contig pair. For each query scaffold/contig, we defined a single block. If two different queries are aligned to a same reference with overlaps, the larger one was selected. All contigs that were aligned to the same reference sequence collinearly were merged to form a larger scaffold. A merging was confirmed if it was supported by short reads alignments. For assembly of PWN rDNA single repeat unit, online searching found three fragments of PWN rDNA published in NCBI GenBank (AB500152.1, AY508034.1, AB294736.1), which have overlaps with each other to assemble a draft rDNA single repeat unit. It includes all expected features such as IGS, 18S, ITS1, 5S, ITS2, and 28S, although 28S was not fully assembled. The assembly gaps were further closed by IMAGE2^49^ using WGS short reads of BxCN. Final BxCN rDNA assembly was examined by aligning short reads to assembled rDNA so that the rDNA assembly sequences could be fixed based on mismatches with short reads. The completeness of the draft genome of BxCN was assessed using Core Eukaryotic Genes Mapping Approach (CEGMA) with a set of core proteins, which are highly conserved ubiquitous proteins in eukaryotic species^50^, to search for orthologs from the new genome. The BxCN genome sequences have been deposited in NCBI (accession number: POCF00000000).

Short reads by whole-genome sequencing of the other 54 pinewood nematode strains were aligned to BxCN assembly, BxCN rDNA and published mitochondrial DNA^51^ using SMALT (http://www.sanger.ac.uk/science/tools/smalt-0) in default settings. For mtDNA and rDNA, a filtration strategy was applied to retain the properly mapped read pairs, which have ≥ 75% of read length mapped, in proper orientations (paired end) and insert sizes (within mean ±3*S.D).

### Construction of subgenomes for phylogenetic analysis

To infer the phylogenetic relationship of sequenced strains with confidence (BxCA was excluded because of the large number of ambiguous alignments caused by genetic diversity), we identified the fragments in the reference genome, the so-called “subgenome”, from which the sequences in each strain were able to be unambiguously identified by short reads alignment. The subgenome in BxCN genome assembly required the minimum coverage of 10×, no InDel, and the sequences with at least 60% read support in all strains. Then, for each strain, sequences in subgenome fragments ≥ 50 bp were concatenated into consensus for further phylogenetic analysis, which covered 14,981,085 bases (18.9% of BxCN assembly) in total and 172,053 (excluding the Ka4 strain) with variations. The subgenomes in mtDNA and rDNA also required a minimum coverage of 10× and no InDel in all strains, but no limit in read support of the sequences and the fragment size. The nucleotides with the highest read support would be selected to build the consensus sequences for further phylogenetic analysis.

Five subgenomes and datasets were constructed: 1) the subgenome of nuclear DNA in 54 strains sequenced in this study and the published strain Ka4; 2) the subgenome of nuclear DNA in 29 DIG strains; 3) the consensus sequences of full-length mtDNA of 55 strains in this study (including BxCA), six from GenBank (**Table S2**) and the Ka4 strain by removing all sites with InDel or “N” in any strain; 4) the subgenome of rDNAs in our 55 strains and Ka4 strain; 5) the consensus sequences of ITS in 55 strains and 112 published ones (**Table S2**), including Ka4. They were used for phylogenetic tree construction and population structure analysis.

### Phylogenetic tree construction and principle components analysis

We applied both Neighbour-joining (NJ) and Maximum likelihood (ML) methods to infer the phylogenetic trees based on the sequences in the subgenomes identified previously in the BxCN assembly, mtDNA and rDNA. The NJ tree was inferred using MEGA 6.0^52^ with the substitution model of maximum composite likelihood, while the ML tree was inferred using RAxML^53^ with the substitution model of “GTRCAT”. The bootstrap analyses of 1000 times were applied in both methods. Population structure was inferred by using a principal component analysis (PCA) based on the single nucleotide polymorphisms (SNPs) with smartpca in the software package of EIGENSOFT^54^. We included all samples in the analysis with default parameters. The top two most informative principal components were used to illuminate the relationship among worldwide strains.

### Functional annotation of protein-coding genes

Gene prediction in the reference genome (BxCN) was inferred using *de novo*-homology- and evidence-based approaches. *De novo* gene prediction was performed on repeat-masked genomes using four gene prediction programs (Augustus, GlimerHMM, SNAP and GeneMark.hmm-ES)^47^. Training gene model sets were generated from subsets of the EST/mRNA and RNA-Seq datasets representing 822 distinct genes of BxCN. To take advantage of the value of the protein-coding gene annotation of the published Japanese PWN genome^28^, we have used a homology-based gene annotation program genBlastG^55^ to find potentially missed gene models in BxCN. RNA-seq reads obtained by sequencing the reverse transcribed mRNA libraries of BxCN were aligned using bowtie^56^. Introns are defined using TopHat^57^.

Functional annotation of proteins encoded by GVs-harboured genes was performed by aligning amino-acid sequences to NCBI invertebrate refseq (v2018-03-12, 1E-50), UniProt (v2017-08-30, 1E-50) and Pfam (v31.0; 1E-5), using BLASTP. Proteins were further classified by Gene Ontology using Blast2GO^58^, the euKaryotic Clusters of Orthologous Groups (KOG)^59^, and KEGG (http://www.genome.jp/tools/kaas), using BLAST (E-value of 1e-5). BUSTED approach^60^ was used to identify positive selection (dN/dS>1, P-value < 0.05). The genomic map was drew using Circos^61^.

## Acknowledgments

This work was supported by the National Key Research and Development Program of China (2016YFC1201100), the National Nature Science Foundation of China (31370501) and the Science and Technology Innovation Program of the Chinese Academy of Agricultural Sciences (CAAS-ASTIP-2017-IVF). Research in the group of N.C. is supported by a Discovery Grant from NSERC Canada. N.C. is also a MSFHR Scholar and a CIHR New Investigator.

## Author contributions

X.C., N.C. and B.X. designed and directed the project. J.Li. performed genome sequences assembled and analysed. R.L. performed gene functional annotation. S.X., X.Yi., X.Z. and J.Lin. participated in bioinformatics analysis. Z.M. prepared DNA samples for sequencing. X.K. provided the map of *Pinus* spp. distributing in China. X.Yan., J. Luo. and F.C. contributed to PWN samples collection, nematode culture and material preparation. Y.L. and L.W. contributed PWN samples. J.Li., X.C., N.C. and B.X. analysed data and wrote the manuscript.

## Supplemental materials

**Fig S1** Synteny between the Chinese BxCN and the published Japanese Ka4 genomes of *Bursaphelenchus xylophilus*.

**Fig S2** PWN strains collected from provinces in China (a) and from the countries in the world (b).

**Fig S3** Phylogenetic relationship of all PWN strains (a) Phylogenetic relationship based on rDNA sequences. (b) Phylogenetic relationship based on mitochondria DNA sequences. Sequence names with * are obtained from genBank.

**Fig S4** Alignment of rDNA sequences between ZJHZ2 and other rDIG strains.

**Fig S5** Phylogenetic relationship based on ITS sequences. (a) ITS sequences from this study only.

(b) ITS sequences from this study and from previously published data.

**Table S1** Sources of PWN strains and information of whole genome sequencing

**Table S2** List of rDNA ITS sequences used in this study from published databases

**Table S3** Pfam annotation of protein-coding genes harboured SNPs with DIG fixation alleles

**Table S4** Protein-coding Genes harboured SNPs with DIG-specific alleles which putatively related to adaptation.

**Table S5** Positive selection genes

## References

1. Webster, J. & Mota, M. Pine wilt disease: global issues, trade and economic impact. In Pine wilt disease: a worldwide threat to forest ecosystems. (eds Mota, M. & Vieira, P.) 1–3 (London: Springer, 2008).

2. Jones, J. T. et al. Top 10 plant-parasitic nematodes in molecular plant pathology. Mol. Plant Pathol. 14, 946–961 (2013).

3. Keller, S. & Taylor, D. R. History, chance and adaptation during biological invasion: Separating stochastic phenotypic evolution from response to selection. Ecology Letters 11, 852–866 (2008).

4. Dwinell, L. D. & Nickle, W. R. An overview of the pinewood nematode ban in North America. Gen. Tech, Rep. SE-55. (US Department of Agriculture Forest Service, Southeastern Forest Experiment Station: Asheville, NC) 13 pp. (1989).

5. Yang, B. J., Wang, Q. L., Zhou, W. D. & Li, Y. C. The resistance of pine species to pine wood nematode, Bursaphelenchus xylophilus. Acta Phytopathologica Sinica 17, 211–214 (1987).

6. Yang, B. J. et al. Studies on the resistance of pine trees to pine wood nematode, Bursaphelenchus xylophilus. Forest Research 6, 249–255 (1993).

7. Wingfield, M. J. & Blanchette, R. A. The pine wood nematode, Bursaphelenchus xylophilus, with stressed trees in Minnesota and Wisconsin: insect associates and transmission studies. Can. J. Forest Research 13, 1068–1076 (1983).

8. Kobayashi, F., Yamave, A. & Ikeda, T. The Japanese pine sawyer beetle as the vector of pine wilt. Ann. Rev. Ent. 29, 115–135 (1984).

9. Kinn, D. N. Incidence of pinewood nematode dauerlarvae and phoretic mites associated with long-horned beetles in central Louislana. Can. J. For. Res. 17, 187–190 (1987).

10. Zhao, L., Niu, H., Fang, G., Zhang, S. & Sun, J. A native fungal symbiont facilitates the prevalence and development of an invasive pathogen-native vector symbiosis. Ecology 94, 2817–2826 (2013).

11. Zhao, B. G., Wang, H. L., Han, S. F. & Han, Z. M. Distribution and pathogenicity of bacteria species carried by Bursaphelenchus xylophilus in China. Nematology 5, 899–906 (2003).

12. Oku, H. Pine wilt toxin, the metabolite of a bacterium associated with a nematode. Naturwissenschaften 67, 198–199 (1980).

13. Cheng, X. Y. et al. Metagenomic analysis of the pinewood nematode microbiome reveals a symbiotic relationship critical for xenobiotics degradation. Scientific Reports 3, 1869 (2013).

14. Cheng, H. R., Lin, M., Li, W. & Fang, Z. The occurrence of a pine wilting disease caused by a nematode found in Nanjing. For. Pest Dis. 4, 1–5 (1983).

15. Song, Y. S. & Zang, X. Q. Analysis of potential geographic distribution of Bursaphelenchus xylophilus in China and quarantine counter measures. For. Pest Dis. 4, 38–41 (1989).

16. Yang, B. J. & Wang, Q. L. Distribution of the pinewood nematode in China and susceptibility of some Chinese and exotic pines to the nematode. Can. J. For. Res. 19, 1527–1530 (1989).

17. Xu, F., Ge, M., Wang, Q., Zhang, P. & Zhu, K. Studies on the masson pine provenances resistance to pine wood nematode disease in China. J. Nanjing For. Univ. 22, 29–33 (1998).

18. Yang, B. J., Pan, H. Y., Tang, J., Wang, Y.Y. & Wang, L. F. Pine wilt disease. Beijing: China Forestry Publishing House; pp 1–263 (2003).

19. Shi, J., Chen, F., Luo, Y.Q., Wang, Z. & Xie, B. Y. First isolation of pine wood nematode from Pinus tabuliformis forests in China. For. Pathol. 43, 59–66 (2013).

20. Lee, C. E. Evolutionary genetics of invasive species. Trends Ecol. Evol. 17, 386–391 (2002).

21. Mallez, S. et al. First insights into the genetic diversity of the pinewood nematode in its native area using New polymorphic microsatellite loci. PLoS One 8, e59165 (2013).

22. Mallez, S. et al. Worldwide invasion routes of the pinewood nematode: What can we infer from population genetics analyses. Biol. Invasions 17, 1199–1213 (2015).

23. Zhou, Z. H., Sakaue, D., Wu, B. Y. & Hogetsu, T. Genetic structure of populations of the pinewood nematode Bursaphelenchus xylophilus, the pathogen of pine wilt disease, between and within pine forests. Phytopathology 97, 304–310 (2007).

24. Jung, J., Han, H., Ryu, S. H. & Kim, W. Microsatellite variation in the pinewood nematode, Bursaphelenchus xylophilus (Steiner and Buhrer) Nickle in South Korea. Genes Genom. 32, 151–158 (2010).

25. Cheng, X. Y., Cheng, F. X., Xu, R. M. & Xie, B.Y. Genetic variation in the invasive process of Bursaphelenchus xylophilus (Aphelenchida: Aphelenchoididae) and its possible spread routes in China. Heredity (Edinb) 100, 356–365 (2008).

26. Vieira, P., Burgermeister, W., Mota, M., Metge, K. & Silva, G. Lack of genetic variation of Bursaphelenchus xylophilus in Portugal revealed by RAPD-PCR analyses. J. Nematol. 39, 118–126 (2007).

27. Fonseca, L. et al. The pinewood nematode, Bursaphelenchus xylophilus, in Madeira Island. Helminthologia 49, 96–103 (2012).

28. Kikuchi, T. et al. Genomic insights into the origin of parasitism in the emerging plant pathogen Bursaphelenchus xylophilus. PLoS Pathog. 7, e1002219 (2011).

29. Palomares-Rius, J. E. et al. Genome-wide variation in the pinewood nematode Bursaphelenchus xylophilus and its relationship with pathogenic traits. BMC Genomics 16, 845 (2015).

30. Sakai, A. K. et al. The population biology of invasive species. Ann Rev Ecol Syst. 32, 305–332 (2001).

31. Pereira, F. et al. New insights into the phylogeny and worldwide dispersion of two closely related nematode species, Bursaphelenchus xylophilus and Bursaphelenchus mucronatus. PLoS One 8, e56288 (2013).

32. Figueiredo, J. et al. Assessment of the geographic origins of pinewood nematode isolates via single nucleotide polymorphism in effector genes. PLoS One 8, e83542 (2013).

33. Yamada, T. & Ito, S. Chemical defense respones of wilt - resistant pine species, Pinus strobes and P. taeda, against Bursaphelenchus xylophilus. Ann. Phytopath. Soc. Japan 59, 666–672 (1993).

34. Stamps, W. T. & Linit, M. J. Interaction of intrinsicand extrinsic chemical cues in the behaviour of Bursaphelenchus xylophilus (Aphelenchida: Aphelenchoididae) in relation to its beetle vectors. Nematology 3, 295–301 (2001).

35. Cardoso, J. C. R., Fellx, R. C., Fonseca, V. G. & Power, D. M. Feeding and the rhodopsin family G-protein coupled receptors in nematodes and arthropods. Front. Endocrin. 3, 157 (2012).

36. Zhang, K. et al. Molecular phylogeny of geographical isolates of Bursaphelenchus xylophilus: implications on the origin and spread of this species in China and worldwide. J. Nematol. 40, 127–137 (2008).

37. Valadas, V., Barbosa, P., Espada, M., Oliverira, S. & Mota, M. The pinewood nematode, Bursaphelenchus xylophilus, in Portugal: possible introductions and spread routes of a serious biological invasion revealed by molecular methods. Nematology 14, 899–911 (2012).

38. Mamiya, Y. History of pine wilt disease in Japan. J. Nematol. 20, 219–226 (1988).

39. Corlett, R. T. Environmental forestry in Hong Kong: 12871-1997. For. Ecol. Man. 116, 93–105 (1999).

40. Tzean, S. S. & Tang, J. S. The occurrence of the pine wood nematode, Bursaphelenchus xylophilus, in Taiwan. Proceedings of the 6th ROC symposium on electron microscopy 1985, 38–39 (1985).

41. Yi, C., Byun, B., Park, J., Yang, S. & Chang, K. First finding of the pinewood nematode, Bursaphelenchus xylophilus(Steiner Buhrer) Nickle and its insect vector in Korea. Res. Rep. For. Res. Ins. Seoul. 38, 141–149 (1989).

42. Mota, M. et al. First report of Bursaphelenchus xylophilus in Portugal and in Europe. Nematology 1, 727–734 (1999).

43. Robertson, L. et al. Incidence of the pinewood nematode Bursaphelenchus xylophlius Steiner & Buhrer. in Spain. Nematology 13, 755–757 (2011).

44. Barrett, R. D. H. & Schluter, D. Adaptation from standing genetic variation. Trends Ecol. Evol. 23, 38–44 (2008).

45. Cheng, X. Y., Xie, P. Z., Cheng, F. X., Xu, R. M. & Xie, B. Y. Competitive displacement of a native species Bursaphelenchus mucronatus by an alien species Bursaphelenchus xylophilus(Aphelenchida: Aphelenchoididae) — a case of successful invasion. Biol. Invasion 11, 205–213 (2009).

46. Li, R. et al. The sequence and de novo assembly of the giant panda genome. Nature 463, 311–317 (2010).

47. Chu, J. S., Baillie, D. L. & Chen, N. Convergent evolution of RFX transcription factors and ciliary genes predated the origin of metazoans. BMC Evol. Biol. 10, 130 (2010).

48. Angiuoli, S. V. & Salzberg, S. L. Mugsy: fast multiple alignment of closely related whole genomes. Bioinformatics 27, 334–342 (2011).

49. Tsai, I. J., Otto, T. D. & Berriman, M. Improving draft assemblies by iterative mapping and assembly of short reads to eliminate gaps. Genome Biol. 11, R41 (2010).

50. Parra, G., Bradnam, K. & Korf, I. CEGMA: a pipeline to accurately annotate core genes in eukaryotic genomes. Bioinformatics 23, 1061–1067 (2007).

51. Sultana, T. et al. Comparative analysis of complete mitochondrial genome sequences confirms independent origins of plant-parasitic nematodes. BMC Evol. Biol. 13, 12 (2013).

52. Tamura, K., Stecher, G., Peterson, D., Filipski, A. & Kumar, S. MEGA6: Molecular Evolutionary Genetics Analysis version 6.0. Mol. Biol. Evol. 30, 2725–2729 (2013).

53. Stamatakis, A. RAxML version 8: a tool for phylogenetic analysis and post-analysis of large phylogenies. Bioinformatics 30, 1312–1313 (2014).

54. Patterson, N., Price, A. L. & Reich, D. Population structure and eigenanalysis. PLoS Genet. 2, e190 (2006).

55. She, R., et al. genBlastG: using BLAST searches to build homologous gene models. Bioinformatics 27, 2141–2143 (2011).

56. Langmead, B. & Salzberg, S. L. Fast gapped-read alignment with Bowtie 2. Nat. Methods 9, 357–359 (2012).

57. Pollier, J., Rombauts, S. & Goossens, A. Analysis of RNA-Seq data with TopHat and Cufflinks for genome-wide expression analysis of jasmonate-treated plants and plant cultures. Methods Mol. Biol. 1011: 305–315 (2013).

58. Conesa, A. et al. Blast2GO: a universal tool for annotation, visualization and analysis in functional genomics research. Bioinformatics 21: 3674–3676 (2005).

59. Koonin, E.V. et al. A comprehensive evolutionary classification of proteins encoded in complete eukaryotic genomes. Genome Biol. 5, R7 (2004).

60. Murrell, B. et al. Gene-wide identification of episodic selection. Mol. Biol. Evol. 32, 1365–1371 (2015).

61. Krzywinski, M. et al. Circos: an information aesthetic for comparative genomics. Genome Res. 19, 1639–1645 (2009).

